# Protein Optimization Evolving Tool (POET) based on Genetic Programming

**DOI:** 10.1101/2022.03.05.483103

**Authors:** Alexander R. Bricco, Iliya Miralavy, Shaowei Bo, Or Perlman, Christian T. Farrar, Michael T. McMahon, Wolfgang Banzhaf, Assaf A. Gilad

## Abstract

Proteins are used by scientists to serve a variety of purposes in clinical practice and laboratory research. To optimize proteins for greater function, a variety of techniques have been developed. For the development of reporter genes used in Magnetic Resonance Imaging (MRI) based on Chemical Exchange Saturation Transfer (CEST), these techniques have encountered a variety of challenges. Here we develop a mechanism of protein optimization using a computational approach known as “genetic programming”. We developed an algorithm called Protein Optimization Evolving Tool (POET). Starting from a small library of literature values, use of this tool allowed us to develop proteins which produce four times more MRI contrast than what was previously state-of-the-art. Next, we used POET to evolve peptides that produced CEST-MRI contrast at large chemical shifts where no other known peptides have previously demonstrated contrast. This demonstrated the ability of POET to evolve new functions in proteins. Interestingly, many of the peptides produced using POET were dramatically different with respect to their sequence and chemical environment than existing CEST producing peptides, and challenge prior understandings of how those peptides function. This suggests that unlike existing algorithms for protein engineering that rely on divergent evolution, POET relies on convergent evolution.

## INTRODUCTION

Natural evolution has produced a myriad of proteins and many of them have been used for medical treatment and recently for diagnostics. But since the beginning of life, natural evolution has only explored a small portion of the protein design space, challenging protein engineers to optimize existing and to even create new protein functions. Directed Evolution is a common and powerful technique to artificially evolve proteins in the laboratory^1^. In general, directed evolution starts from a template protein that has a function similar to the desired one. Next, a library of mutant proteins is generated often by using error-prone DNA polymerase and screened for the ‘fittest’ protein that shows the most desired feature. This first generation will then serve as a template for the next generation, and the procedure is repeated until a suitable protein with respect to the particular feature is found (**Fig. 1a**).

**Figure 1.**
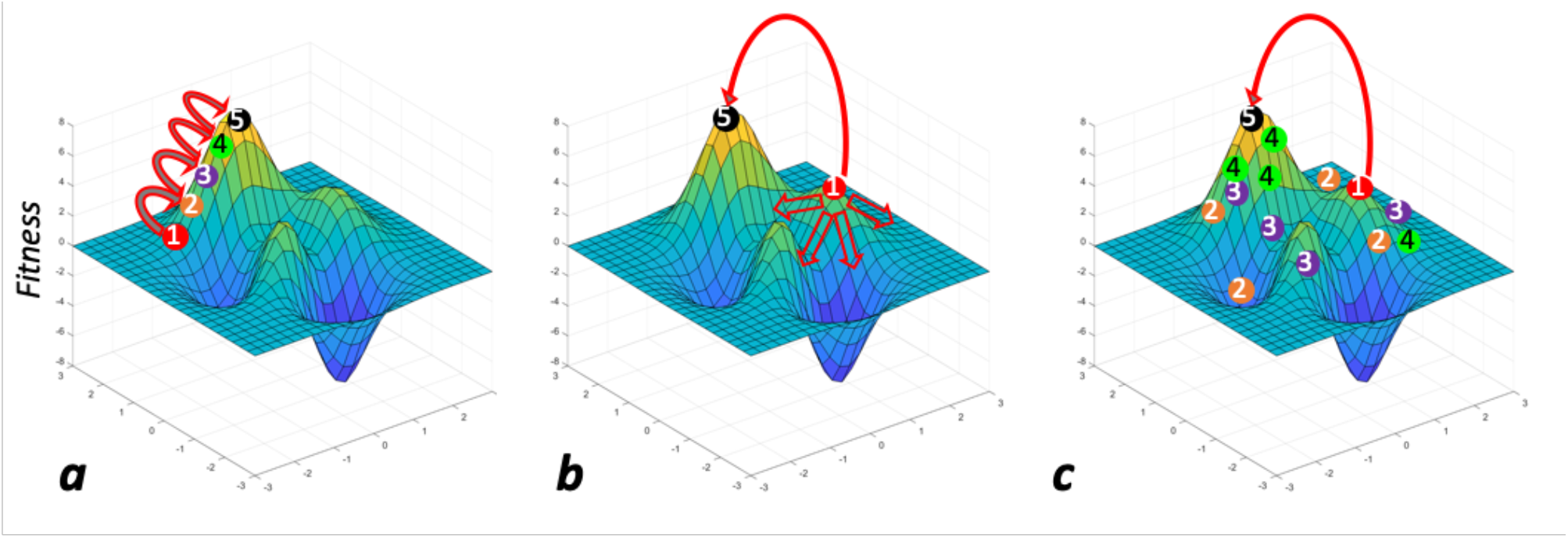
The principles of POET. (**a**) illustration of conventional directed evolution, where in each evolution cycle one mutant exhibit better fitness and thus, used as a template for the following generation of evolution. (**b**) Often, in directed evolution, the protein fitness reaches a local maximum and consequently, all the mutants exhibit lower fitness (empty arrows). In this case it is impossible to predict which mutant should be used as a template to achieve improved fitness (the route from ① to ⑤). (**c**) In the case of POET, the route from① to ⑤ is not determined by stepwise mutagenesis and adhering to the parental protein, but rather by generating libraries of peptides that cover broadly all the search space. Each generation helps to shape a set of rules that determine the next set of peptides. This way all the search space of the fitness landscape is covered and consequently minimizing the probability of missing the absolute maximum.

Despite its effectiveness, directed evolution comes with several limitations. For many proteins, the experimental evaluation process is very time consuming. Many of the mutants produce silent mutations which do not carry on to later generations. Furthermore, optimizing proteins requires navigation through a complex fitness landscape, with optimization trajectories often leading to a dead-end, unless several mutations occur at once^2, 3^ (**Fig. 1b**). Deploying a novel *Protein Optimization Evolving Tool (POET)* based on genetic programming can make it possible to overcome these challenges by exploring a wider range of the protein design space (**Fig. 3c**). POET relies on the principle of convergent evolution, i.e., when species/proteins have different origins but have developed similar features. This is in contrast to divergent evolution, in which separate species evolve differently from a common origin. Thus, POET allows for the identification of new peptides/proteins with desired features that could not have been discovered with any of the traditional protein engineering tools. POET utilizes all the search space; and even protein variants that do not show improvement over previous generations to provide useful information that can lead to improvement of the next generation. Hence, the POET algorithm is potentially a game changer protein design tool that can be implemented into numerous protein engineering applications.

*Evolutionary Computation* is a field in computer science, studying algorithms inspired by biological evolution. *Genetic Programming (GP)*^4, 5^ is among powerful evolutionary computation techniques that evolves solutions to difficult structural design tasks as a general problem solver. In the context of protein engineering, GP was used to predict trans-membrane domains and omega loops in proteins^4^, to evolve energy functions for evaluating protein structures^6^, and more recently, to predict protein-protein interactions related to disease^7^. This earlier work demonstrates the capability of GP to model features in the protein search domain, and in particular its ability to extract features relevant for a prediction task. This is a central capability in biological applications where often high-dimensional inhomogeneous datasets are used as input to predict output values. In addition to creating predictive models, the underlying mechanisms of GP allows it to come up with novel models, often on first sight surprising or even counter-intuitive to the user^8^. Over the last decades, GP has proven to produce human-competitive solutions to many problems^9^.

To evaluate the potential of POET to evolve ultra-sensitive proteins and peptides, we decided to focus on solving the problem of sensitivity of a specific class of peptide-based probes used for magnetic resonance imaging (MRI) of molecular targets. The peptides can be detected with MRI via a contrast mechanism, termed Chemical Exchange Saturation Transfer (CEST). CEST is based on the dynamic exchange process between an exchangeable proton (hydrogen atom) and the surrounding water protons^10, 11^. While this contrast was demonstrated to be most efficient for poly-L-lysine by van Zijl and colleagues^12^ and later on was optimized for several other peptides^13, 14^, contrast improvement using rational protein design remained a challenge. Moreover, when the peptides were genetically encoded and expressed in cells or rodents, the background contrast that was generated by cellular metabolites was very high^15-19^. Therefore, there was a need to generate peptides that provide MRI contrast at a frequency where the background CEST contrast is low. This in evolutionary terms, will be creating a new function. To achieve both goals --improving existing contrast and creating a new contrast - we deployed POET to evolve peptides that provide high CEST contrast in a frequency dependent manner.

## MATERIALS AND METHODS

### Peptide synthesis and preparation

The peptides determined by POET, were obtained from Genscript (Piscataway, NJ). Each peptide was dissolved to a concentration of 5 mg/mL in deionized water. To keep differences in pH from interfering with the CEST effect, each peptide was titrated to a pH of 7.2 using 0.1 M HCl and 0.1 M NaOH.

### MRI parameters

In earlier generations MRI data acquisition was obtained using a vertical bore 11.7 Tesla Bruker Avance system with the 0.2 mL samples placed in the imaging coil and kept at 37°C during imaging. The first scan is a WASSR scan used to determine the exact frequency of water in the sample so that it may be adjusted accordingly^20^. The second scan is a CEST scan made from a modified RARE sequence (TR/effective TE = 10000/4.5 ms, RARE factor = 32, FOV = 17 × 17 mm^2^, slice thickness = 1.2 mm, matrix size = 64 × 64, spatial resolution = 0.27 × 0.27 mm^2^) including a continuous-wave saturation pulse of 4 s, saturation powers of 1.2 μT, 2.4 μT, 3.6 μT, 4.7 μT, 6.0 μT, 7.2 μT, 10.8 μT, 12.0 μT covering saturation frequencies from -10 to +10 ppm offset from water in steps of 0.27 ppm. Starting with the 5^th^ generation we acquired MRI data using a horizontal bore 7T Bruker preclinical MRI. The samples were placed within an imaging phantom custom designed and produced by 3D printing specifically for this task. Each group of samples is run through two scans. The first scan is a WASSR scan used to determine the exact frequency of water in the sample so that it may be adjusted accordingly^20^. The second scan is a CEST scan made from a modified RARE sequence, with a RARE factor of 16, and a TR of 10000 ms. Saturation pulses were applied as a block pulse for 4000 ms, and a saturation power of 4.7 μT covering saturation frequencies from -7 to 7 ppm offset from water in steps of 0.2 ppm. Each generation was scanned multiple times to ensure accuracy. Data processing was done with an in-house MATLAB script^21^. Finally, the amide proton chemical exchange rate was measured by Quantitation of Exchange with Saturation Power (QUESP)^22^ (McMahon MT, et al, Magn Reson Med 2006;55:836-847) using an ultra-fast Z-spectroscopy method ^23^ on a 14 Tesla Bruker NMR spectrometer where the saturation power was varied from 0.2 to 14.8 μT. The saturation time was 5 s, the TR was 10 s, the number of averages was 8, and the temperature was maintained at 37°C.

### Exchange rate calculation and simulation

The amide proton exchange rate was quantified by Bloch-McConnell equation fitting of the power dependence of the 14T amide proton signal using custom written software (MATLAB). The amide proton signal was extracted from the ultra-fast Z-spectrum by 4-pool (water, amide, amine and hydroxyl proton pools) Lorentzian fitting {Zaiss, 2011 #1243}Some of the peptides displayed poor fits, likely due to a range of amide proton exchange rates being present in the peptides. Improved fits of the QUESP data were obtained by fitting the low and high saturation power data separately.

To investigate the relationship between the amide proton exchange rate and the asymmetric magnetization transfer ratio MTR_asym_ at different saturation pulse powers, a simulation study was performed based on the numerical solution of the Bloch-McConnell equations implemented in MATLAB (MathWorks)^24^. The parameters used were longitudinal water relaxation (T_1_) of 1600 ms for both the water and solute pools, transverse relaxation time (T_2_) of 50 ms and 1 ms for the water and solute pools, respectively, and a solute concentration of 200 mM with a chemical shift of 5 ppm. The simulated acquisition protocol used an echo time (TE) of 20 ms, a repetition time (TR) of 15 s, a continuous saturation pulse of 5 s, applied at 9 to -9 ppm frequency offsets, with 0.25 ppm intervals, and a readout flip angle of 90º, under a 7T main magnetic field (B_0_). The examined exchange rates varied uniformly between 100 to 2000 Hz with 1 Hz increments.

### Genetic Programming

POET algorithm is a multi-platform GP tool written in the Python programming language. The computational experiments were run on Michigan State University’s High-Performance Computing Center (HPCC) systems. Each POET replicate uses only a single CPU core (2.5 GHz) and 8 gigabytes of RAM. At each generation of the experiment, 100 replicates of POET are executed in parallel using different random seeds to evolve protein-function models able to predict the CEST contrast of peptide sequences. These replicates allow POET to explore various regions of the ‘search space at the same time to find fitter models. After the evolution of these models, the fittest one of them in each generation is employed for predicting new optimized peptides with respect to their CEST contrast levels. To do so, a population of 10,000 random peptide sequences is initialized and evolved by applying an iterative evolutionary algorithm. In this algorithm, each of the sequences in each iteration undergoes point mutation and is evaluated using the fittest previously evolved POET model. If a mutation is not beneficial then it is reverted, and the sequence will move to the next population unchanged. Otherwise, if the mutation is beneficial, the change is applied to the sequence to be added to the next population. This process is performed for an arbitrary number of iterations (usually set to 1000) until fitter predicted peptide sequences are found. The top 10 predicted peptide sequences are chosen to be tested in the lab. Following lab measurements, these predicted peptides are added to the POET’s training dataset, increasing POET’s chances to learn more meaningful motifs in the next generation of the experiment and enabling it to predict fitter and more optimized proteins in the future. At the very start, 42 data points (peptide sequences and their respective CEST contrast values) were available in the POET training dataset. Furthermore, in each generation of the experiment, approximately 10 new predicted peptides were added to the dataset after wet-lab measurements. In the final generation of the experiment, 128 data points were available causing each execution of POET to take up to 35 hours to evolve fit sequence-function models.

Detailed explanations of the computational aspects of POET can be found in a sister article ^25^.

## RESULTS AND DISCUSSION

### Developing Protein Optimization Evolving Tool (POET) based on Genetic Programming

Genetic Programming much like many other evolutionary algorithms follows the basic principles of evolution. A population of random solutions to a given problem is generated as the first generation. The fitness of each of these solutions is evaluated and quantified as a measurement for their performance. The solutions with the highest fitness values are more likely to be selected to create the next generation of solutions after being impacted by evolutionary operators such as crossovers and mutations. Crossover is a reproduction mechanism analogous to sexual reproduction. Usually in crossover two parent solutions are selected to create two new offspring. A common way to do so is to combine genetic codes for each of the parent solutions in a manner that the offspring will contain parts from both parents but is not identical to either of them. Mutation usually occurs after crossover and has a chance to randomly modify a small detail of solutions. The general goal of GP is to evolve solutions to reach a specified fitness level. In other words, to find a solution that satisfies the solving criteria of a problem.

As a first step of developing POET, we incorporated GP to evolve CEST predicting models represented by tables of motifs and weights. Motifs are recurring patterns in protein sequences and their respective weight represents the impact of that pattern in calculating the CEST contrast of a given protein. For example, a motif could be Glycine-Arginine-Arginine (GRR) or Arginine-Lysine (RK) and their initial weights could be –0.60 and 4.39 units, respectively (**Fig. 2a, b and Table S-1**). POET models attempt to find their motifs in given protein sequences and add the weights of the found motifs to generate a score value correlated with the CEST contrast of that protein. POET attempts to find and evolve models that best predict the CEST contrast. POET generates an initial random population of 100 models which can have up to 50 rows of motifs and weights. Evaluation of these models is done by comparing the score values from these models with the actual CEST contrast levels of proteins in a training dataset. These models are then compared by how well they can predict the CEST contrast measured from the training data (**Fig. 2c**).

**Fig. 2.**
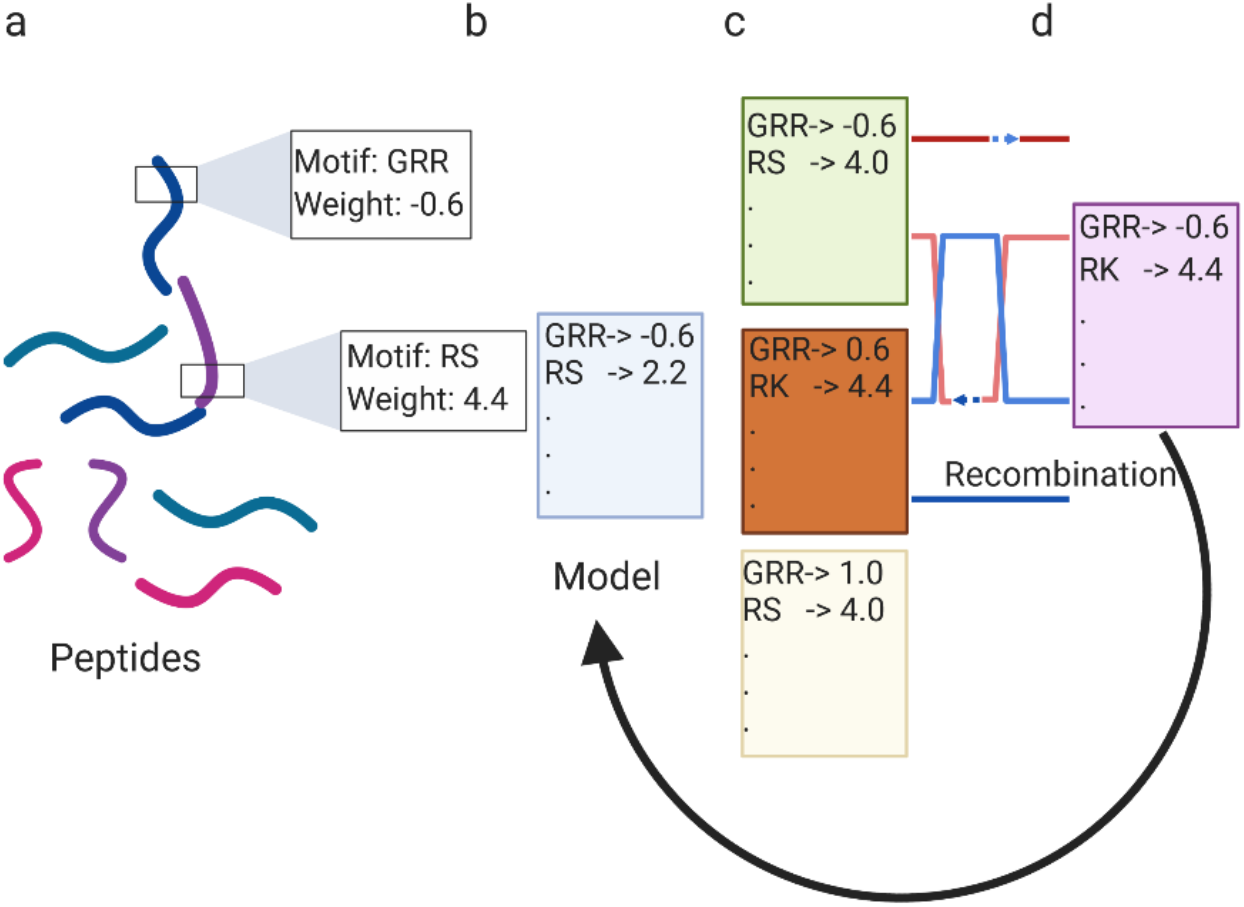
Schematic illustration of POET. (a-d) paradigm and workflow.

POET uses a selection mechanism called the *Tournament Selection* to choose the parent models from the population (**Fig. 2c, d**). Five models are selected, and their fitness values are compared against each other. The two models with the highest fitness are selected to reproduce two offspring models. POET divides the table of each parent model into two sections of A and B and then generates two offspring models, each of which will contain section A of one parent table and section B of the other. POET uses mutational operators that modify the weights and motifs of the model tables. Across 5,000 to 50,000 iterations of this algorithm (**Fig 2. arrow**), the motif-weight pairs that are most important and accurate at predicting the training dataset are maintained, and those that are poor at improving the training dataset are discarded, causing the model to develop in an analogous manner to Darwinian evolution^25^.

### POET develops a library of contrast producing peptides

Much like in evolution, the fittest peptide can be either selected from a large population (in this case training set) or alternatively, many generations can compensate for a smaller population.

The evolution using POET was performed for ten generations, and the resulting contrast relative to K12 (a sequence of 12 lysines), can be seen in **Fig. 3c**. K12 was chosen as a peptide for comparison due to the high contrast it produces and similarities to other reported results from Poly-L-Lysine^12, 14, 16^. For each generation we have obtained a library of ten synthetic peptides which were termed for convenience “*CESTides*”. **Fig. 3a, b** shows z-spectra (CEST-spectra) and MTR_aysm_ plots respectively. The amplitude of the peak of the plot at 3.6 ppm - the amide resonance frequency was used to generate the generational plot **Fig. 3c**. As can be seen in **Fig. 3**, within 10 generations, POET generated a CESTide that displays a 4-fold increase in the MRI contrast. Interestingly, the best CESTide was produced in generation 7.

**Figure 3:**
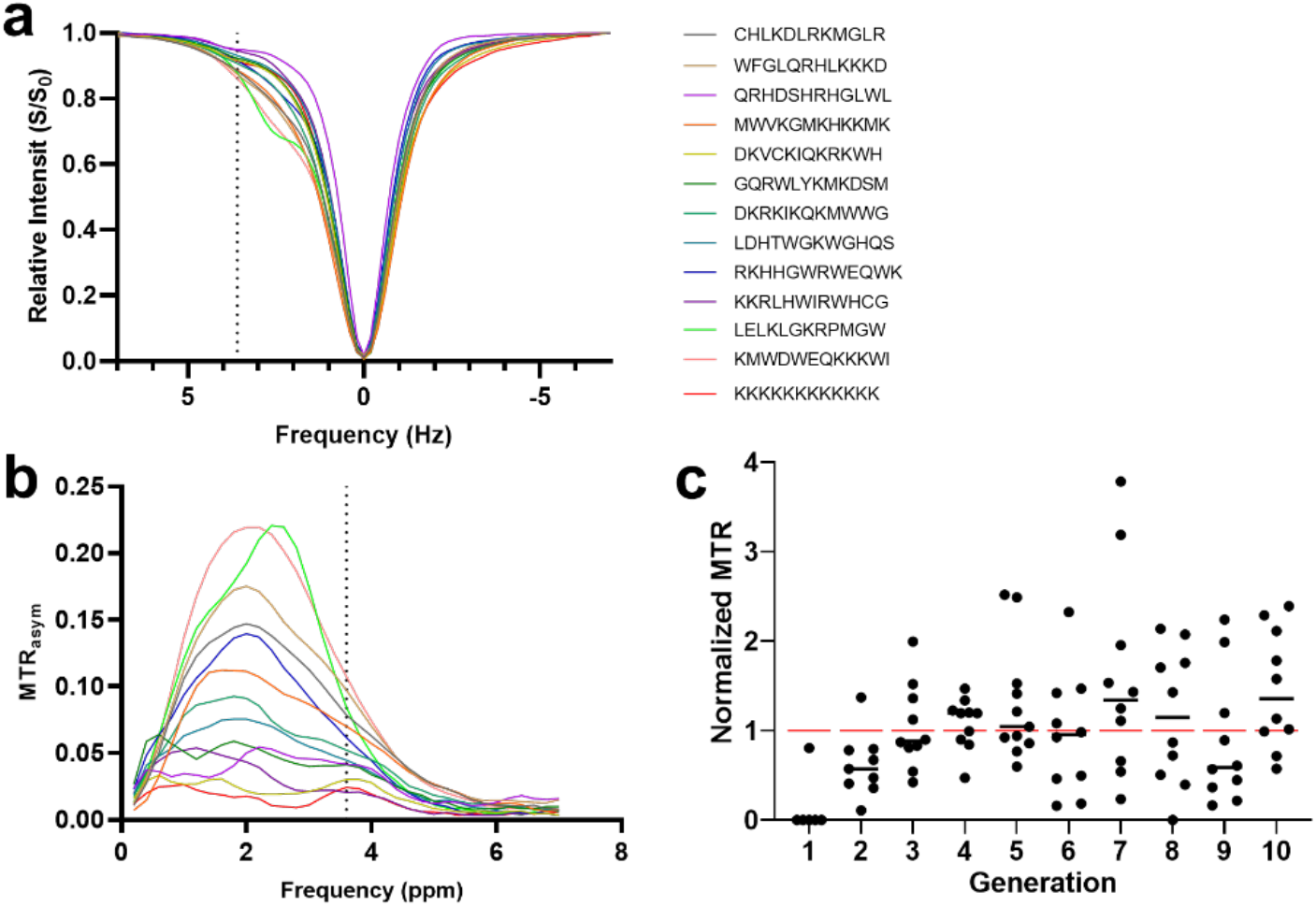
Improvement of CESTides by POET. (**a**) Z-spectra collected from all the peptides in the 5^th^ generation of protein optimization using POET. (**b**) Resulting CEST contrast from these peptides. (**c**) The MTR_asym_ is normalized against the contrast generated by K12 in the same experiment to provide a consistent comparison across experiments and plotted with respect to the generations.

### Sequence diversity of CESTides

POET was able to generate a large variety of different chemistries (**Fig. S-1**), many of which wouldn’t be discovered by directed evolution on a feasible time scale. Traditionally, the general convention is that peptides that are suitable for generating CEST contrast should be positively charged^26, 27^. However, our findings demonstrate that (**Fig. 4 and Fig. S-2)** good CESTides can deviate from the poly-L-lysine like sequence. This is especially important for designing a new version of genetically encoded CEST based reporters ^19^. Less charged reporters reduce intracellular interactions with other proteins while the use of more varied amino acids increases the intracellular expression level of the reporter as it is not dependent on the supply of a single amino acid. Moreover, the diversity in the CESTides isoelectric point (pI) can allow tailoring the reporter to different cellular environments.

**Figure 4:**
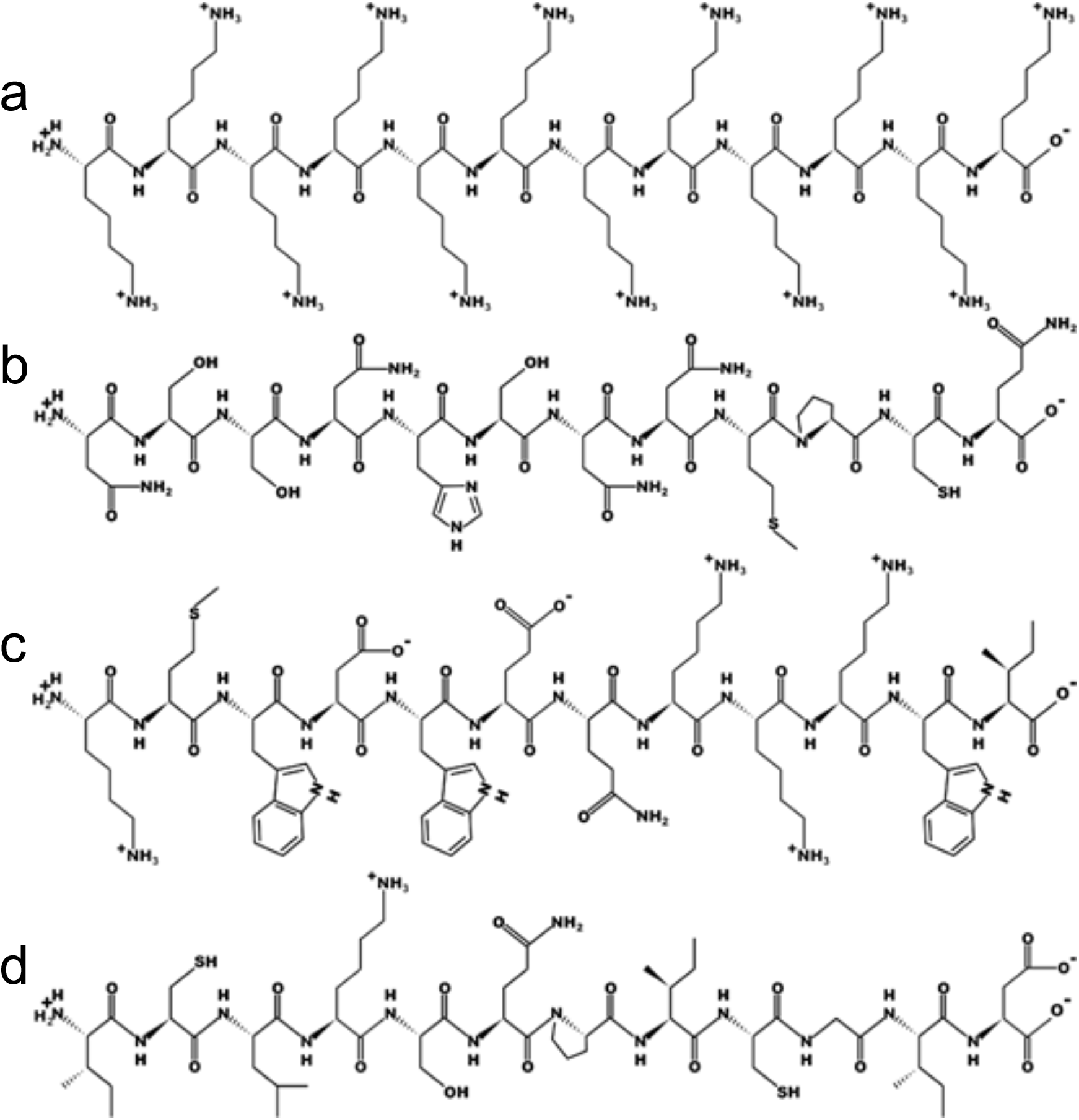
Structure of four representative distinct peptides. **(a)** K12; KKKKKKKKKKKK; Theoretical pI/Mw: 11.04 / 1556.10. **(b)** A peptide from generation 2 has a neutral pI, yet generates contrast higher than the K12; NSSNHSNNMPCQ; Theoretical pI/Mw: 6.73 / 1332.38. **(c)** a peptide from generation 5 that generates contrast that is approximately 4 times larger than K12: KMWDWEQKKKWI; Theoretical pI/Mw: 9.53 / 1706.04. **(d)** a peptide from generation 7 that generates contrast that is twice that of K12 but has an acidic pI: ICLKSQPICGID

### Exchange rate calculations

There are three factors that determine the optimal CEST contrast (MTR_asym_); the chemical shift of the exchangeable proton (Δω), the saturation power (ω_1_) and the optimal exchange rate (k_ex_). While simulations can predict what the optimal three factors are, only Δω and ω_1_ are easy to control experimentally. In contrast, the exchange rate is completely dependent on the chemical formulation of the contrast agent^28^. Hence, it is complicated to predict *in silico* the k_ex_ for a contrast agent with a single exchangeable proton^29, 30^ and even harder to do so for a peptide with at least ten exchangeable protons^22^. We used computational simulations to examine what the optimal exchange rate would be. Our simulations show that peptides with exchangeable amide protons at 3.6 ppm, the k_ex_ that provides the maximal MTR_asym_ should be around 1473 Hz, while peptides with exchangeable protons at 5.0 ppm the maximal MTR_asym_ values are achieved at around 1497 Hz. Therefore, we aimed with POET to evolve peptides with k_ex_ as close as possible to these values. Since k_ex_ is an absolute number that is dependent only on the peptide sequence and structure as well as on the chemical environment of the peptide (i.e., pH, temperature etc.,) and is independent of the field strength, we determined the k_ex_ for selected peptides using a 14 Tesla MR spectrometer, which provides better spectral resolution. **Table 1** shows improvement in the exchange rate of peptides with evolution. The measured exchange rates are of course an average of the exchange rates of all the exchangeable protons with the same chemical shift. We note that the best fits of the 14 Tesla QUESP peptide data were obtained by fitting the low and high saturation power data separately to quantify the slow and fast exchange rate components, respectively Remarkably, two peptides, KYTKTRKQSSKA and NSSNHSNNMPCQ showed average k_ex_ that are 1.78 and 1.94 times faster than K12. Therefore, using POET we were able to optimize the proton exchange rate of selected peptides through evolution and consequently improve the CEST contrast.

**Table 1.** Normalized CEST Contrast and exchange rate for selected peptides.

| Peptide<br>Sequence | Generation | Normalized CEST Contrast (%)<sup>1</sup> | | | $k_{ex}$ (Hz) |
| --- | --- | --- | --- | --- | --- |
|  |  | Amide | Amine | OH | Amide |
| KKKKKKKKKKKK<sup>2</sup> | 0 | 2.18 | 0.04 | 0.0 | 423 |
| KPWHGCASRTKR | 4 | 4.28 | 4.95 | 6.41 | 548 |
| DKVCKIQKRKWH | 5 | 2.91 | 1.74 | 0.0 | 422 |
| KKRLHWIRWHCG | 5 | 2.27 | 4.67 | 0.0 | 195 |
| CCWHNPKWRRTR | 3 | 2.05 | 5.23 | 7.19 | 433 |
| KYTKTRKQSSKA | 3 | 5.42 | 2.19 | 4.71 | 754 |
| NSSNHSNNMPCQ | 2 | 5.26 | 0.91 | 1.40 | 822 |

<sup>1</sup> normalized to 1 mM peptide concentration, $B_1=6 \mu\text{Tesla}$
<sup>2</sup> K12

### Learning by POET

We sought to examine the differences between the peptides generated by POET to determine if POET was converging toward a solution. This was calculated via the nearest neighbor distance from peptides in the same generation using Grantham distance, which takes into consideration differences between the size, charge, and hydrophobicity of different amino acids^31^. The basic assumption is that amino acids that are similar in chemical composition, polarity and molecular volume are more likely to change throughout evolution as they are less disruptive to protein function. To determine whether POET was learning and converging on a solution, we compared the Grantham distance between the peptides discovered with POET with peptides that were generated randomly. We first examined the intergenerational nearest neighbor distance (**Fig. 5a**), by comparing finding the shortest Grantham distance within each peptide’s generation and all prior generations. As the Grantham distance decreased with an increase in the number of generations, this implies that learning took place since it shows that the predictions of POET are more similar than would be generated by randomness and are decreasing in distance faster. Next, we examined the intragenerational nearest neighbor distance by comparing each peptide to all peptides in the same generation to determine the most similar peptide (**Fig 5b**). We find that the distance stays lower than the random simulation, implying that there is a form of selection occurring since the distance is lower than that of random peptides. The distance is not decreasing by generation which suggests that POET is not converging on a solution, which would show the predictions decreasing in distance as they all approach the same global maximum.

**Figure 5.**
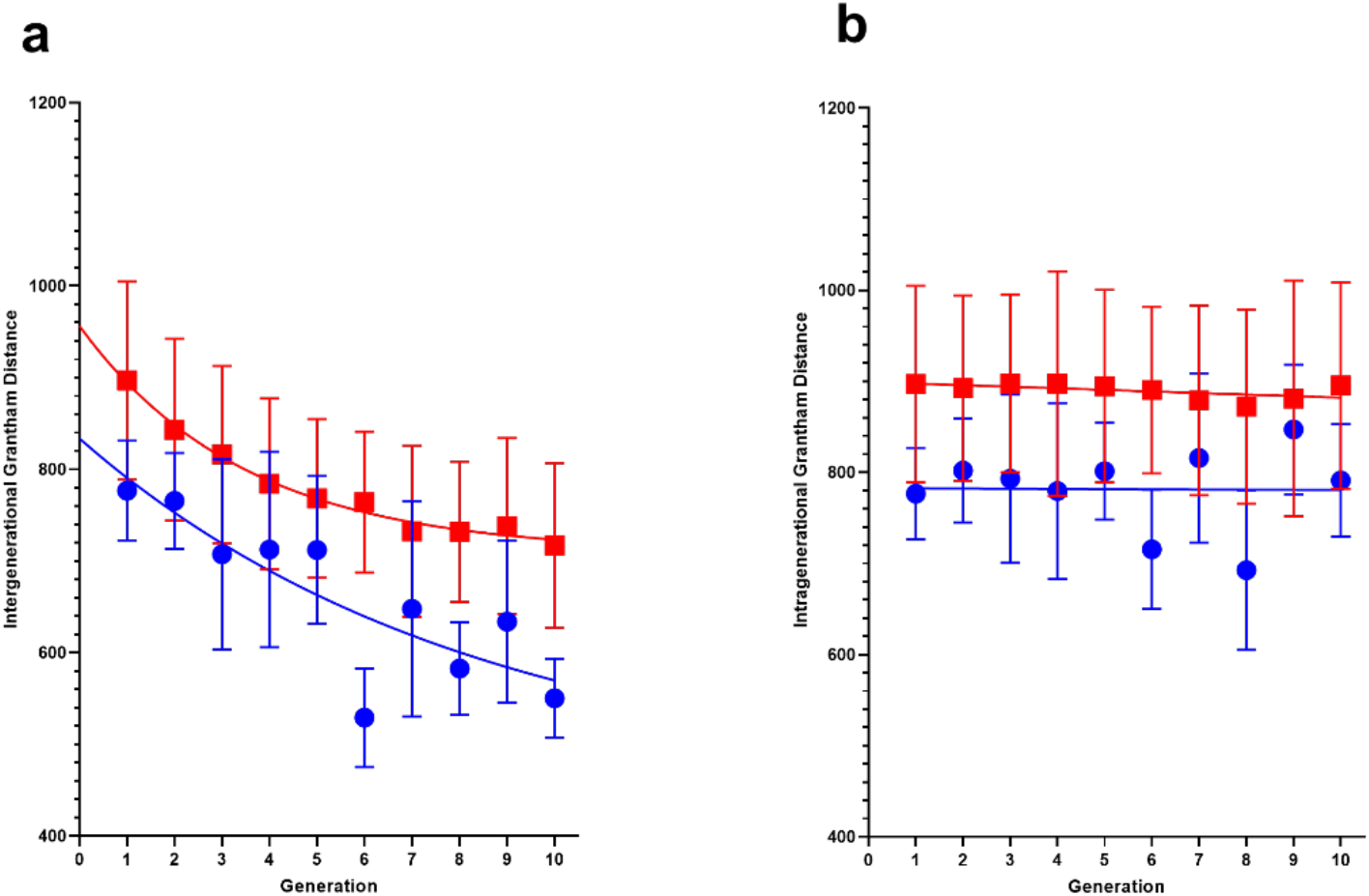
Grantham distance between discovered CESTides. **(a)** Intergenerational distance, where peptides are compared to those in their own generation and all prior ones. **(b)** Intragenerational distance, where peptides are only compared to those in the same generation. The peptides discovered using POET are blue circles (mean ± 95% confidence interval, simulated peptides generated randomly, are shown as red squares (mean ± 95% CI). Each dataset has a trendline fit to an exponential decay curve (a), or linearly (b).

### Evolving new function at 5 ppm

So far, we have deployed POET to improve the existing function of a peptide, that is the CEST contrast provided at a chemical shift of 3.6 ppm. Next, we wanted to test the potential of POET to create a new function that could not be achieved based on our current knowledge or by computational simulations. We have chosen to evolve peptides that generate CEST contrast at a chemical shift of 5 ppm downfield from the water resonance frequency. The rational was that there is relatively little background CEST signal from endogenous peptides and cellular metabolites at that chemical shift, resulting in a higher contrast to noise ratio (CNR). Additional advantage of agents that exchange farther downfield is that they allow higher k_ex_ resulting in better contrast^28, 32, 33^. Since we acquired the CEST spectra for the libraries described above from -7 ppm to 7 ppm, we were able to use the data as a training library for evolving an optimized CESTide generating contrast at a chemical shift of 5 ppm. The development of contrast at 5 ppm can be seen in **Fig. 6**. We have evolved 23 CESTides that generate contrast at 5 ppm that is greater than the contrast obtained at the same frequency from the training data. The MTR_asym_ in **Fig. 6** is normalized to the contrast that K12 produces at 3.6 ppm (i.e., K12 at 3.6 ppm produces MTR_asym_ = 1). Moreover, POET evolved five new CESTides that produce CEST contrast at 5 ppm that is between 0.6-0.8 of the contrast that K12 produces at 3.6 ppm. Considering the low CEST background that cells and tissue produce at 5 ppm^30, 34^, it is anticipated that assembling these peptides will result in a reporter gene that is more sensitive than the 3.6 ppm based genetically encoded CEST reporters^15-19^. These findings indicate that POET can assist in mining the fitness landscape to identify peptides with a new function that cannot evolve otherwise.

**Figure 6:**
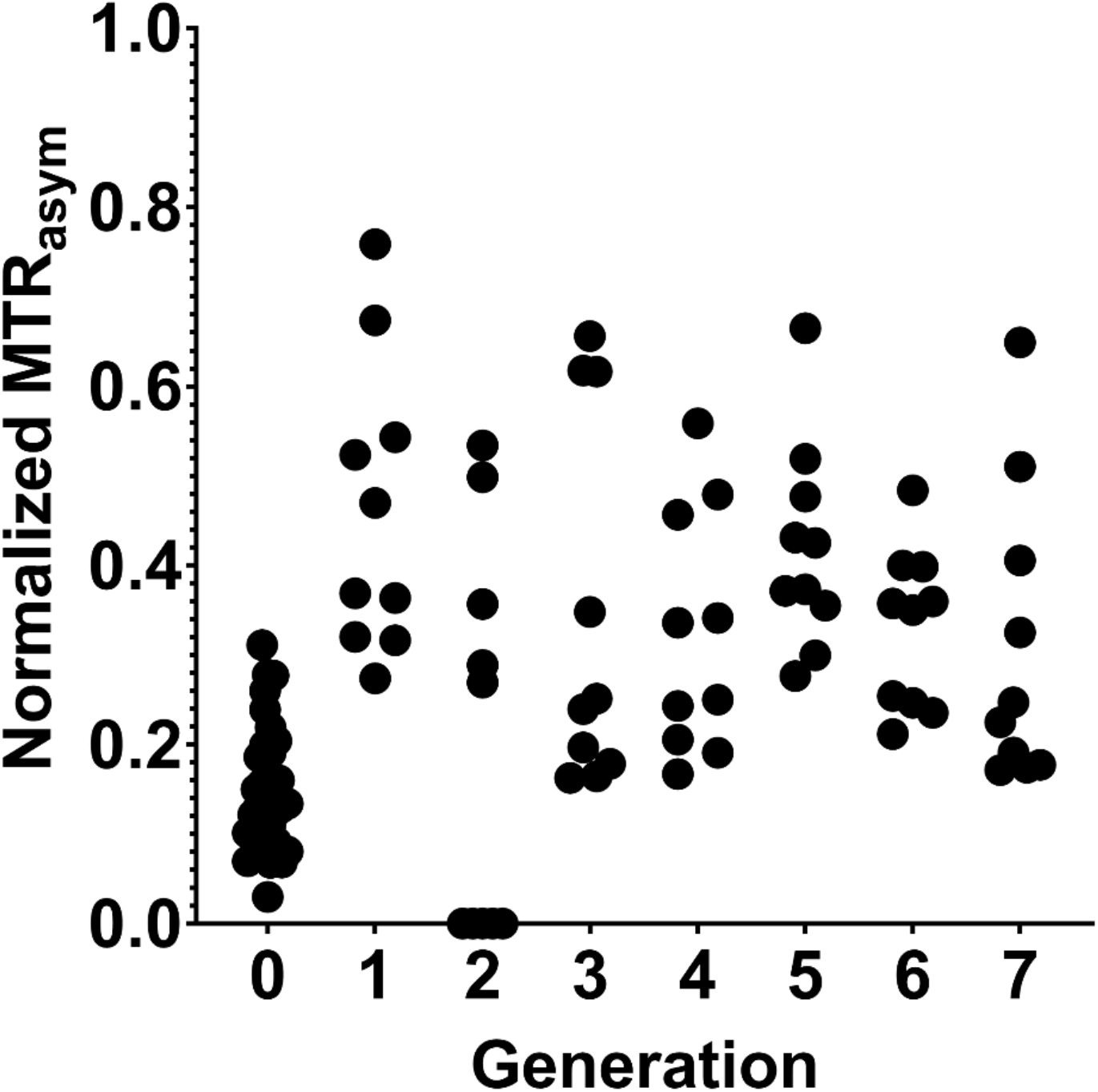
Contrast generated at 5 ppm. Each generation of peptides developed by using POET at 5 ppm. The MTR_asym_ is normalized against the contrast generated by K12 at 3.6 ppm in the same experiment to provide a consistent comparison across experiments.

In recent years, CEST has been used for measuring *in vivo* temperature changes,^35^ pH,^36, 37^ enzyme activity^38, 39^, metal ions^40^, metabolites^41^, glycogen and glucose^42, 43^, glutamate, glycoproteins,^44^ and glycosaminoglycan^45^. Recently, CEST MRI has been performed in the beating heart to detect fibrosis after myocardial infarction in mice^46^ and for *in vivo* mapping of creatine kinase metabolism^47^. We have previously demonstrated that CEST can be used to monitor sustained drug release,^48^ and to sense cellular signaling using a genetically encoded biosensor^49^. Moreover, we have repeatedly demonstrated that reporter genes based on CEST MRI can be used to monitor gene expression in a 3D cell culture^27, 50^ *in vivo* in rodents^30^ or in a live pig heart^51^. In many of these examples, the CEST contrast is generated from a unique exchangeable proton. In this case, to improve the CEST contrast it is sufficient to optimize the exchange rate of this unique proton and this could be done using rational design^27^. However, when designing a peptide for imaging, with multiple protons that exchange in different resonance frequencies and different rates, the optimization is too complex and is beyond the current rational design capabilities. Thus, using tools like POET that combines machine learning algorithms and evolutionary principles with experimental measurement, is ideal for peptide optimization.

We do note that some of the POET optimized peptides (e.g. KKRLHWIRWHCG) have lower amide exchange rates relative to K12. However, the amine contrast at 2 ppm was significantly greater for these peptides indicating that the increased MTR_asym_ at 3.6 ppm for these optimized peptides has contributions also from the amine exchangeable protons. This can be seen in Figure 3b where a strong amine MTR_asym_ is observed at 2 ppm for some of the optimized peptides. Thus, POET can optimize and exploit both amide and amine exchangeable protons to maximize the MTR_asym_ at 3.6 ppm.

## CONCLUSIONS

Here we demonstrated that POET can be used to evolve peptides to produce substantially more CEST contrast than PLL after only a few generations. Furthermore, POET was successfully deployed to evolve peptides with new function, i.e., reporting MRI contrast from peptides at a resonance frequency that could not be detected otherwise. POET generated CESTides could potentially be assembled into the next generation of MRI reporter gene^52^ with improved sensitivity over previous generations of reporters. Since POET requires only a small set of input peptide sequences and their corresponding biological quantitively measured function to evolve models that predict better peptide function, it is anticipated that POET can be used for evolving of peptides in numerous applications.

## ACKNOWLEDGEMENTS

The authors acknowledge financial support from the NIH/NINDS: R01-NS098231; R01-NS104306 NIH/NIBIB: R01-EB031008; R01-EB030565; R01-EB031936; P41-EB024495 NIH/NIDDK: R01-DK121847, NIH S10-OD023406 and NSF 2027113. We thank Rita Martin for help in editing the manuscript and Dr. Jorden Schossau for the helpful discussions.

