## SUPPLEMENTAL MATERIALS for "Protein Optimization Evolving Tool (POET) based on Genetic Programming"

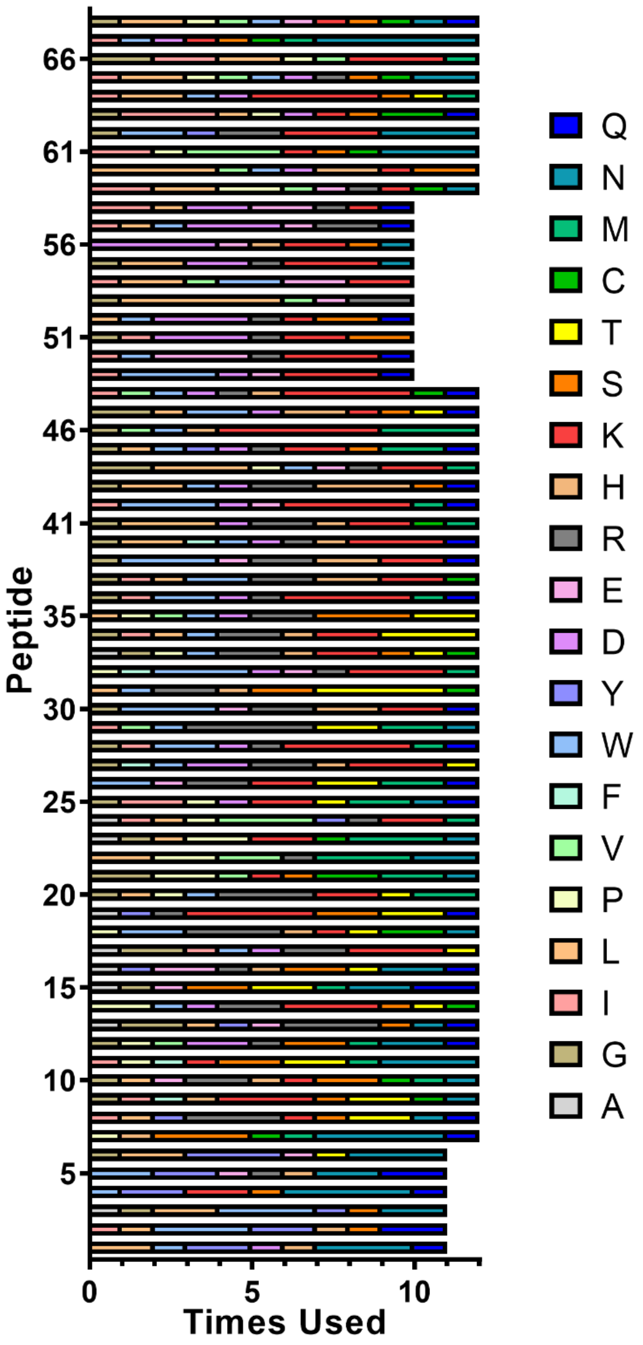


**Figure S-1: Amino Acid Composition of Different Peptides.** Each peptide that was examined is shown to see its different amino acid composition.

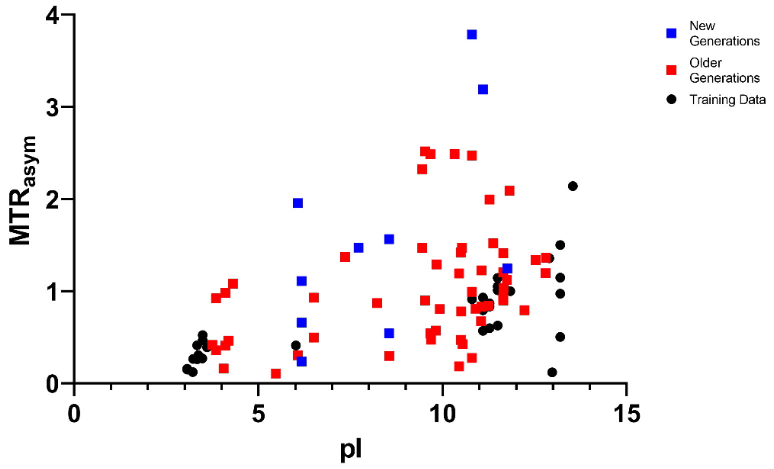


**Figure S-2: Peptide $MTR_{asym}$ with respect to the pI .**

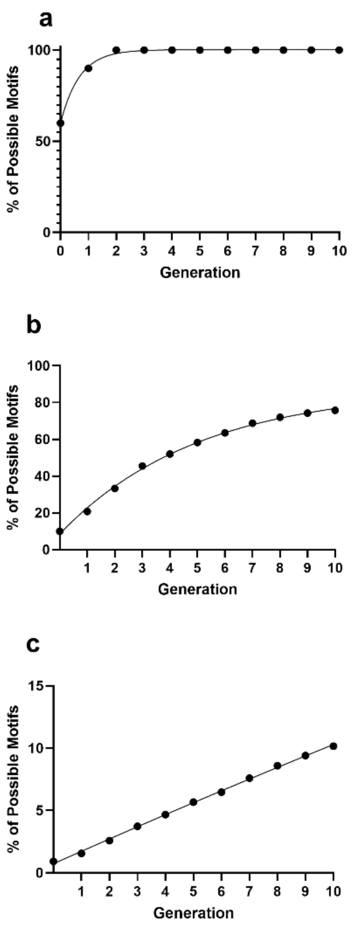


**Figure S-3: Search Space Representation.** The figures show the number of possible motifs for POET to learn from the data set it has been provided. The motifs are divided based on their length, for 1 amino acid long motifs (a), 2 amino acid long motifs (b), and 3 amino acid long motifs (c).

**Table S-1: Example POEM table**. This table lists the motifs and weights that were used in the first generation of POET evolutions.

| **Motif** | **Weight** |
|---|---|
| GRR | -0.60 |
| RRS | 1.65 |
| RS | 2.04 |
| HR | 0.83 |
| RT | 3.54 |
| GR | 2.83 |
| RK | 4.39 |
| KK | 1.89 |
| NK | 1.65 |
| RP | 0.07 |
| R | 1.65 |
| D | -0.29 |
| K | 1.81 |
| E | -0.58 |
| T | 0.73 |
